# Invasive Chinese tallow is associated with distinct rhizosphere microbial communities, core microbiomes, and functional potential relative to native Loblolly pine

**DOI:** 10.64898/2026.10.09.758132

**Authors:** Maruf Khan, Bishrant Pant, Ahmad H. Kabir

**Affiliations:** Department of Biology, Lamar University, Beaumont, TX 77705, United States

**Keywords:** Core microbiome, Environmental filtering, Forests, Invasive plant, Rhizosphere microbiome, Soil microbial networks

## Abstract

Plant invasions frequently reshape belowground microbial communities, yet their effects on rhizosphere microbiome structure and function remain poorly understood. We compared the rhizosphere bacterial and fungal microbiomes of invasive Chinese tallow (*Triadica sebifera*) and native Loblolly pine (*Pinus taeda*) using 16S rRNA gene and ITS amplicon sequencing, together with microbial, functional, and soil analyses. Chinese tallow exhibited significantly greater bacterial and fungal richness and distinct microbial community composition than Loblolly pine. The invasive rhizosphere was enriched in plant-beneficial bacterial taxa, including *Streptomyces, Bradyrhizobium*, and *Rhizobium*, and ectomycorrhizal-associated fungi such as *Russula* and *Cenococcum*. This enrichment is consistent with enhanced nutrient acquisition, plant growth promotion, and stress tolerance, potentially favoring Chinese tallow establishment and persistence. Further, core microbiome and co-occurrence network analyses revealed extensive reorganization of persistent (*Streptomyces, Russula*) and highly connected microbial taxa that may be associated with Chinese tallow invasion. Functional predictions indicated greater representation of carbohydrate degradation, nitrogen fixation, plant growth promotion, nutrient mineralization, and ectomycorrhizal guilds in the Chinese tallow rhizosphere, whereas lignin degradation and saprotrophic or pathogen-associated fungal guilds were relatively more abundant under Loblolly pine. Soil physicochemical properties were also strongly correlated with dominant bacterial and fungal taxa, indicating that host-mediated microbial recruitment and environmental filtering jointly shaped rhizosphere community assembly. Overall, our findings identify rhizosphere microbiome reassembly as an important belowground feature associated with Chinese tallow invasion.

## 1. Introduction

Biological invasions are among the leading drivers of global biodiversity loss and ecosystem alteration, profoundly influencing community structure, nutrient cycling, and ecosystem functioning. Invasive plant species frequently modify belowground processes through changes in litter quality, root exudation, and nutrient acquisition strategies, thereby reshaping rhizosphere microbial communities by selectively recruiting beneficial microorganisms and altering microbial interactions (Coats & Rumpho, 2014; Meng et al., 2025). As rhizosphere microorganisms are essential for nutrient cycling, decomposition, and plant health, invasion-induced microbial shifts may generate positive plant–soil feedbacks that promote invasion success (Wang et al., 2026; Lowry et al., 2024). Understanding how invasive plants influence belowground microbial communities is therefore essential for clarifying the ecological processes associated with invasion success (Bardgett & van der Putten, 2014).

The rhizosphere microbiome is composed of diverse bacterial and fungal communities that interact closely with plant roots and perform essential ecological functions. Bacteria promote nitrogen fixation, nutrient mobilization, and pathogen suppression, whereas fungi contribute to nutrient acquisition, mycorrhizal associations, and organic matter decomposition (Han et al., 2026; Brar et al., 2024). Bacteria such as *Bradyrhizobium, Rhizobium*, and *Streptomyces* contribute to biological nitrogen fixation, nutrient mobilization, disease suppression, and organic matter degradation, whereas ectomycorrhizal fungi including *Russula* and *Cenococcum* improve nutrient acquisition and plant stress tolerance (Compant et al., 2010; Bonito et al., 2014; Trivedi et al., 2020). Plant species strongly influence rhizosphere microbial assembly through host-driven selection, resulting in host-specific microbial communities with distinct functional capabilities (Hodgson et al., 2025; Yue et al., 2023). Advances in high-throughput sequencing have greatly improved our understanding of rhizosphere microbiomes by enabling comprehensive analyses of microbial diversity, community dynamics, core microbiomes, co-occurrence networks, and predicted ecological functions (Philippot et al., 2013; Trivedi et al., 2020).

Invasive plants frequently modify soil properties, including pH, nutrient availability, and organic matter, thereby indirectly reshaping soil microbial communities and reinforcing invasion success (Trivedi et al., 2020; Fierer & Jackson, 2006). Chinese tallow (*Triadica sebifera* L.) is one of the most aggressive woody invasive species in the southeastern United States, where it rapidly colonizes forests, and disturbed habitats (Wang et al., 2012), often displacing native vegetation such as Loblolly pine (*Pinus taeda* L.). Chinese tallow exhibits rapid growth, prolific seed production, high phenotypic plasticity, and broad environmental tolerance, enabling it to establish dense monospecific stands across diverse habitats in the southeastern United States (Rout & Callaway, 2012). Previous studies have shown that Chinese tallow accelerates litter decomposition, alters soil nutrient availability, and reshapes soil microbial communities, potentially facilitating invasion success (Yang et al., 2014; Montez et al., 2021).

Despite increasing evidence that Chinese tallow modifies soil microbial communities, little is known about how bacterial and fungal microbiomes respond simultaneously, how host-specific core microbiomes and microbial interaction networks are reorganized, and whether these community shifts are accompanied by changes in predicted microbial functions. To address these knowledge gaps, we compared the rhizosphere bacterial and fungal microbiomes associated with native Loblolly pine and invasive Chinese tallow using 16S rRNA gene and ITS amplicon sequencing. We integrated analyses of microbial community dynamics, co-occurrence networks, core microbiomes, predicted functional potential, and correlations with soil physicochemical properties to provide a comprehensive assessment of host-associated microbial assembly. This study provides new insights into microbial patterns associated with the ecological success of one of the most problematic invasive tree species in North America.

## 2. Materials and methods

### 2.1. Study site and sample collection

Rhizosphere soil samples were collected from native Loblolly pine (*Pinus taeda* L.) at Dujay Sanctuary (30.33° N, 94.40° W), Hardin County, Texas, USA, and from invasive Chinese tallow (*Triadica sebifera* L.) at Ninth Avenue, Port Arthur (29.93° N, 93.95° W), Jefferson County, Texas, USA. The Hardin County site (30.33° N, 94.40° W) lies within the Piney Woods ecoregion, where soils are sandy to loamy, and well to moderately well drained. These soils are mainly derived from coastal plain sediments and are commonly classified as Ultisols and Spodosols, reflecting advanced weathering, low base saturation, and relatively low natural fertility. In contrast, the Jefferson County site (29.93° N, 93.95° W) is situated closer to the Gulf Coast and is characterized by finer-textured alluvial and coastal plain soils with higher soil moisture, and greater organic matter accumulation (NRCS, 2026; NRCS, 2008). The dominant soil orders include Alfisols, Entisols, and Histosols, which are strongly influenced by coastal hydrology and periodic flooding. The physicochemical properties of rhizosphere soils from the two sampling locations are provided in Supplementary Table S1.

For sample collection, three healthy, mature trees of each species separated by at least 10 m were selected as biological replicates. Surface litter was carefully removed before sampling. Soil was excavated around the base of each tree to a depth of approximately 15 cm using a sterile shovel to expose fine roots. Root systems were gently lifted with the surrounding soil, and loosely attached bulk soil was removed by gentle shaking. Soil that remained tightly adhered to fine roots was collected using sterile spatulas, designated as rhizosphere soil, placed into sterile polypropylene tubes, transported to the laboratory on ice, and stored at −80°C until DNA extraction.

### 2.2. DNA extractions and amplicon sequencing

Rhizosphere soil DNA was extracted using a modified cetyltrimethylammonium bromide (CTAB) method (Clarke, 2009). Approximately 0.2 g of rhizosphere soil was transferred into sterile microcentrifuge tubes, and CTAB extraction buffer was added to lyse microbial cells and release nucleic acids. Samples were thoroughly vortexed and incubated at 65 °C to enhance cell disruption and DNA recovery. An equal volume of chloroform: isoamyl alcohol (24:1, v/v) was then added, mixed gently, and centrifuged to separate the aqueous phase. The upper aqueous phase containing DNA was carefully transferred to a new sterile tube, and DNA was precipitated using cold isopropanol. The DNA pellet was washed with 70% ethanol, air-dried, and dissolved in TE buffer. Extracted DNA was treated with RNase A and stored at −20 °C until further analysis. DNA quality and concentration were assessed using a NanoDrop spectrophotometer (Fisher Scientific, USA).

Rhizosphere microbial communities were characterized using Illumina amplicon sequencing by targeting the bacterial 16S rRNA gene and the fungal internal transcribed spacer (ITS) region. Amplicon libraries were prepared by amplifying the bacterial 16S rRNA (V3V4) gene using the primer pair 338F (5′- ACTCCTACGGGAGGCAGCAG-3′) and 806R (5′- GGACTACHVGGGTWTCTAAT-3′), and the fungal internal transcribed spacer (ITS) region using the primer pair ITS1FI2 (5′- GAACCWGCGGARGGATCA-3′) and ITS2 (5′- GCTGCGTTCTTCATCGATGC-3′). Paired-end sequencing (2 × 250 bp) was performed on the Illumina NovaSeq 6000 platform.

### 2.3. Data processing and microbial community analysis

Raw sequence reads were quality-filtered and adapter-trimmed using Cutadapt (Martin, 2011), followed by denoising, chimera removal, and inference of amplicon sequence variants (ASVs) using the DADA2 pipeline (Callahan et al., 2016). Reads assigned to chloroplast and mitochondrial DNA were excluded from the bacterial dataset prior to downstream analyses. Taxonomic assignment of bacterial ASVs (amplicon sequence variants) was performed against the SILVA 138 reference database (Quast et al., 2013), whereas fungal ASVs were classified using the UNITE database (Nilsson et al., 2019). All downstream analyses were performed in R using the phyloseq (McMurdie & Holmes, 2013), and related packages. Alpha diversity was estimated using the Observed ASVs, Chao1 richness, Shannon diversity, and Simpson diversity indices after rarefaction to an even sequencing depth. Differences in alpha diversity between Loblolly pine and Chinese tallow rhizosphere communities were evaluated using Student’s *t*-tests. Beta diversity was assessed using Principal Coordinates Analysis (PCoA) based on Bray– Curtis dissimilarity matrices calculated from ASV abundance data. Community differences between treatments were tested using permutational multivariate analysis of variance (PERMANOVA).

### 2.4. Microbial co-occurrence network analysis

Microbial co-occurrence networks were constructed separately for bacterial and fungal communities using ASV-level relative abundance data. ASVs with mean relative abundance <0.1% were removed to reduce sparsity. Pairwise Spearman correlations were calculated among the remaining ASVs using the Hmisc package (Harrell, 2012). Only correlations with |r| ≥ 0.6 and P < 0.05 were retained to construct undirected networks with igraph (Pechlivanis et al., 2024). Hub ASVs were identified using centrality metrics, including degree, betweenness, closeness, and eigenvector centrality. The 20 ASVs with the highest degree centrality were selected as hubs and taxonomically annotated at the genus level where possible.

### 2.5. Core microbiome analysis

Core microbiome analysis was performed separately for bacterial and fungal communities at the genus level. ASVs were agglomerated to the genus level, and relative abundances were calculated by transforming sequence counts to proportional abundances within each sample. For each treatment group, the prevalence and mean relative abundance of each genus were determined. Core genera were defined as those detected in at least 70% of samples within a treatment (prevalence ≥70%) and having a mean relative abundance of at least 0.1%. The mean relative abundance of core genera was calculated across biological replicates, and the ten most abundant core genera were visualized as stacked bar plots for comparison between native Loblolly pine and invasive Chinese tallow rhizosphere communities.

### 2.6. Functional prediction analysis

The functional potential of bacterial and fungal communities was inferred from taxonomic composition. For bacteria, ASVs were agglomerated to the genus level, and predicted ecological functions were assigned using the Functional Annotation of Prokaryotic Taxa (FAPROTAX) database by matching bacterial genera to curated functional categories (Louca et al., 2016). Functional abundances were summarized for each sample and averaged across biological replicates within each treatment. For fungi, ecological guilds were assigned using the FUNGuild database based on ITS taxonomic classifications (Nguyen et al., 2016). The relative abundance of each fungal guild was calculated for each sample and averaged across treatments. The 10 most abundant bacterial functional groups and fungal guilds were visualized as horizontal bar plots to compare the predicted functional profiles of rhizosphere microbial communities associated with native Loblolly pine and invasive Chinese tallow.

### 2.7. Correlation analysis between soil physicochemical properties and microbial communities

Relationships between soil physicochemical properties and dominant microbial taxa were evaluated separately for bacterial and fungal communities in R. ASVs were agglomerated to the genus level, sequence counts were transformed to relative abundances, and the 20 most abundant genera were selected for correlation analysis. Pairwise associations between genus-level relative abundances and measured soil physicochemical properties were assessed using Spearman’s rank correlation coefficients. The statistical significance of each correlation was determined using Spearman’s correlation tests, and correlations with *P* < 0.05 were considered significant. Correlation matrices were visualized as clustered heatmaps using the pheatmap package, with hierarchical clustering applied to both microbial genera and soil variables to identify groups exhibiting similar correlation patterns. Significant correlations were indicated by asterisks in the heatmaps.

## 3. Results

### 3.1. Bacterial and fungal community diversity and composition

Principal Coordinates Analysis (PCoA) based on Bray–Curtis dissimilarity revealed distinct clustering of bacterial and fungal rhizosphere communities associated with native Loblolly pine and invasive Chinese tallow (Figure 1A, C). For the bacterial community, the first two principal coordinate axes explained 43.1% and 23.3% of the total variation, respectively. For the fungal community, the first two axes accounted for 42.7% and 20.9% of the variation, respectively (Figure 1A, C). Alpha diversity analyses showed differences in microbial richness between the two rhizosphere communities (Figure 1B, D). For bacterial communities, the Observed ASVs and Chao1 richness indices were significantly higher in the Chinese tallow rhizosphere than in the Loblolly pine rhizosphere (*P* < 0.05), whereas Shannon and Simpson diversity indices did not differ significantly between the two groups (*P* > 0.05). Similarly, fungal communities associated with Chinese tallow exhibited significantly greater Observed ASVs and Chao1 richness than those associated with Loblolly pine (*P* < 0.05), while no significant differences were detected for Shannon or Simpson diversity indices (*P* > 0.05) (Figure 1B, D).

**Figure 1.**
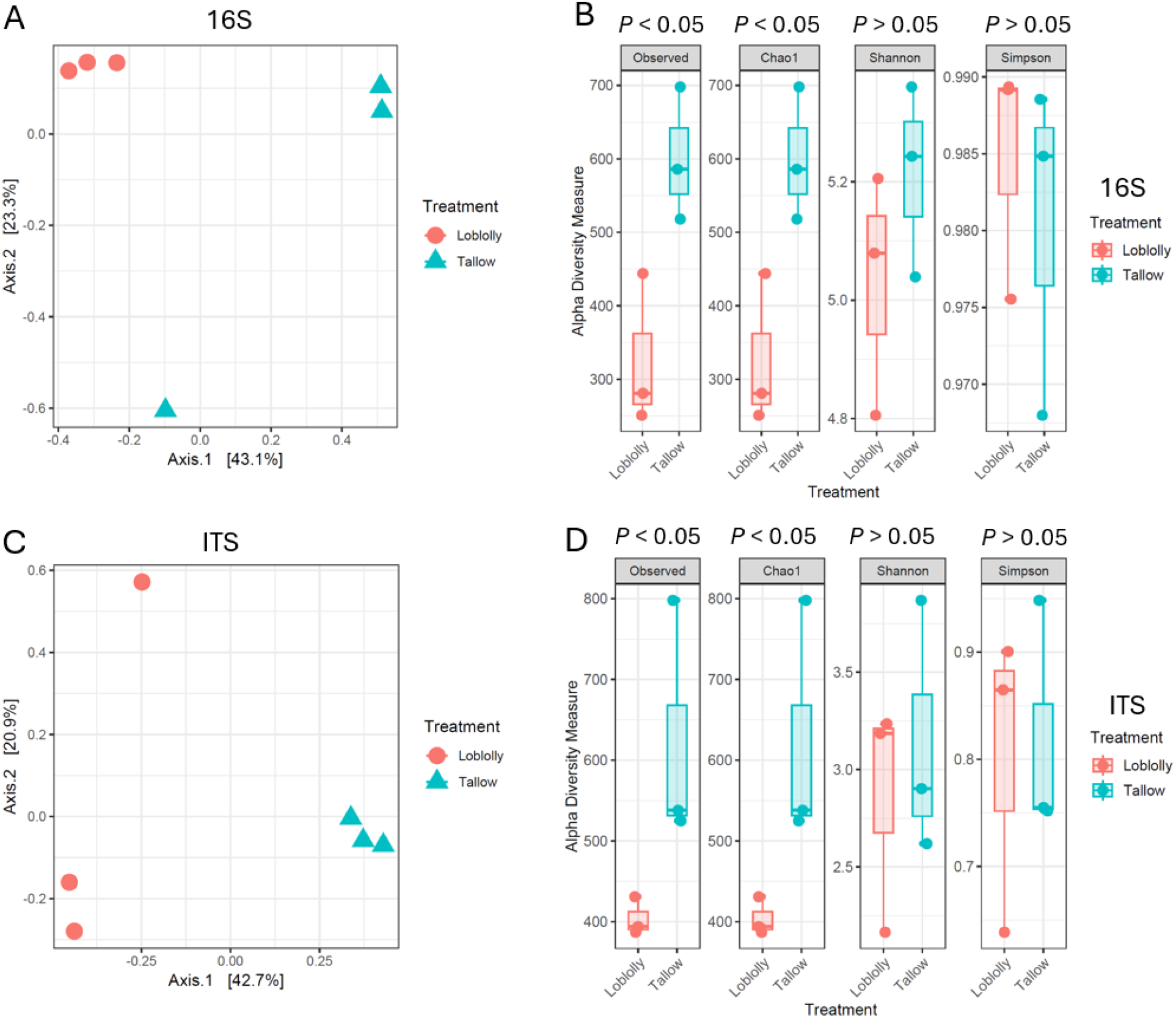
Bacterial and fungal community diversity and composition associated with native Loblolly pine and invasive Chinese tallow rhizospheres. (A, C) Principal Coordinates Analysis (PCoA) showing differences in bacterial and fungal community composition between rhizosphere samples collected from loblolly pine and Chinese tallow. (B, D) Alpha diversity metrics including Observed ASVs (amplicon sequence variant), Chao1 richness, Shannon diversity, and Simpson diversity indices for Loblolly pine and Chinese tallow rhizosphere bacterial and fungal communities. Boxplots show the median, interquartile range, and data distribution, with individual points representing biological replicates.

### 3.2. Relative abundance of dominant bacterial taxa

The composition of the dominant bacterial communities differed between the rhizospheres of native Loblolly pine and invasive Chinese tallow (Figure 2). At the family level, Streptomycetaceae, Micromonosporaceae, Mycobacteriaceae, and Lactobacillaceae exhibited significantly greater relative abundances in the Chinese tallow rhizosphere than in the Loblolly pine rhizosphere (*P* < 0.05), whereas Acetobacteraceae was significantly more abundant in the Loblolly pine rhizosphere (Figure 2A). At the genus level, the dominant bacterial taxa included *Streptomyces, Rhizobium, Occallatibacter, Nocardia, Mycobacterium, Latilactobacillus, Haliangium, Conexibacter, Bradyrhizobium, Aeromyrmexbacter, Actinospica, Actinoplanes, Acidothermus*, and *Acidibacter* (Figure 2B). Among these, *Streptomyces* showed a significantly higher relative abundance in the Chinese tallow rhizosphere, whereas *Latilactobacillus* and *Mycobacterium* were significantly higher (*P* < 0.05) in the Loblolly pine rhizosphere compared to Chinese tallow (Figure 2B).

**Figure 2.**
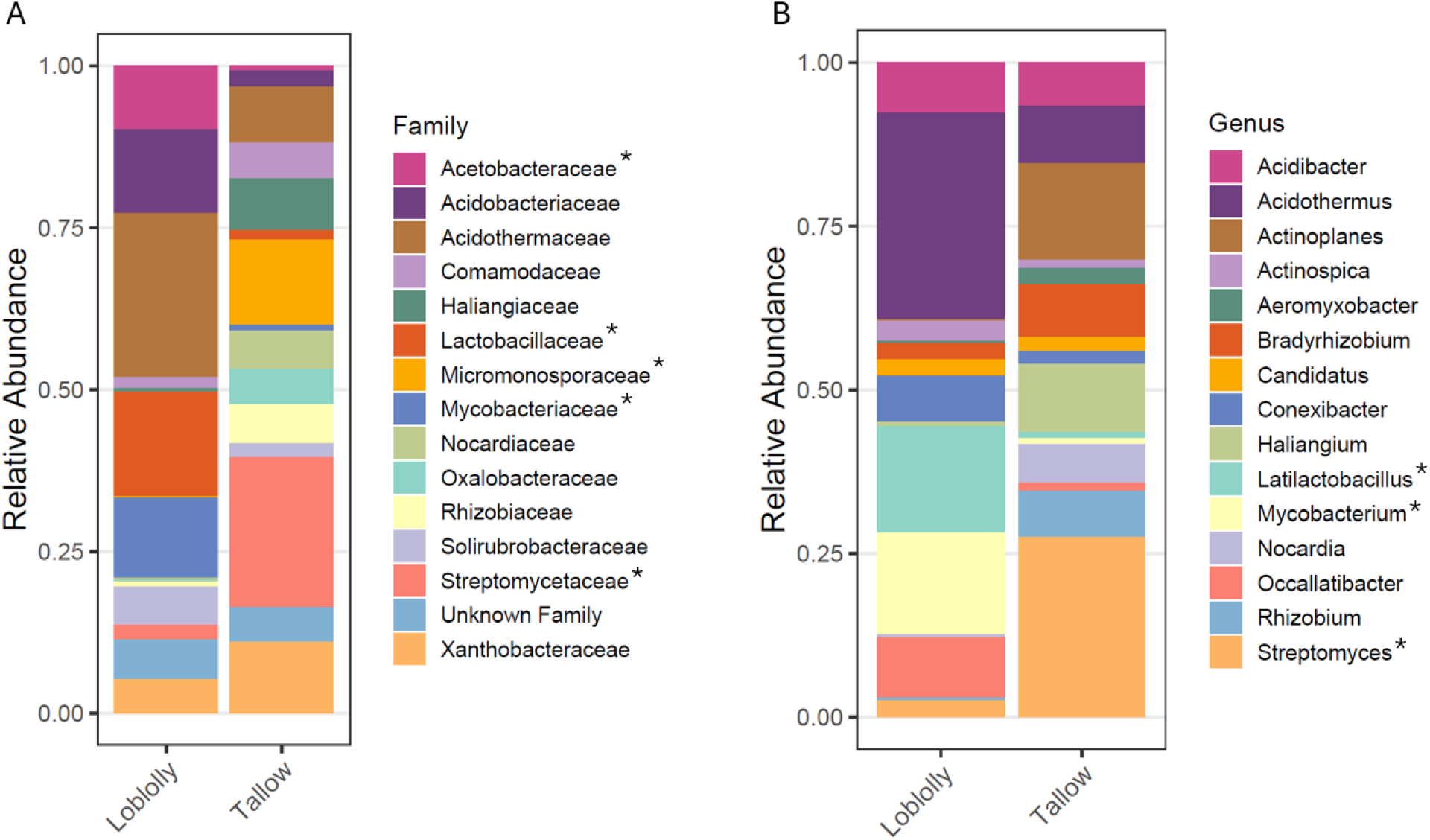
Relative abundance of the top 15 bacterial families (A) and genera (B) associated with Loblolly pine and Chinese tallow rhizospheres based on 16S rRNA gene amplicon sequencing. Asterisks (*) denote taxa that differed significantly in relative abundance between Loblolly pin and Chinese tallow rhizospheres (*P* < 0.05). Each treatment consisted of three independent biological replicates.

The relative abundances of the most abundant fungal families and genera also differed between the rhizosphere communities of native Loblolly pine and invasive Chinese tallow (Figure 3). At the family level, Glomeraceae, Gloniaceae, Massariaceae, Nectriaceae, and Sclerodermataceae exhibited significantly greater relative abundances in the Chinese tallow rhizosphere than in the Loblolly pine rhizosphere (*P* < 0.05). In contrast, Dermateaceae, Helotiaceae, Russulaceae, and Thelephoraceae were significantly more abundant in the Loblolly pine rhizosphere (Figure 3A). At the genus level, the dominant fungal taxa included *Atractiella, Cenococcum, Cladophialophora, Diaporthe, Fusarium, Hymenoscyphus, Penicillium, Pezicula, Russula, Scleroderma, Scutellinia, Stagonospora, Talaromyces*, and *Thelephora* (Figure 3B). Among these, *Russula, Cenococcum, Scleroderma, Scutellinia*, and *Stagonospora* showed greater relative abundances in the Chinese tallow rhizosphere, whereas *Fusarium, Hymenoscyphus, Pezicula*, and *Thelephora* were relatively more abundant in the Loblolly pine rhizosphere (Figure 3B).

**Figure 3.**
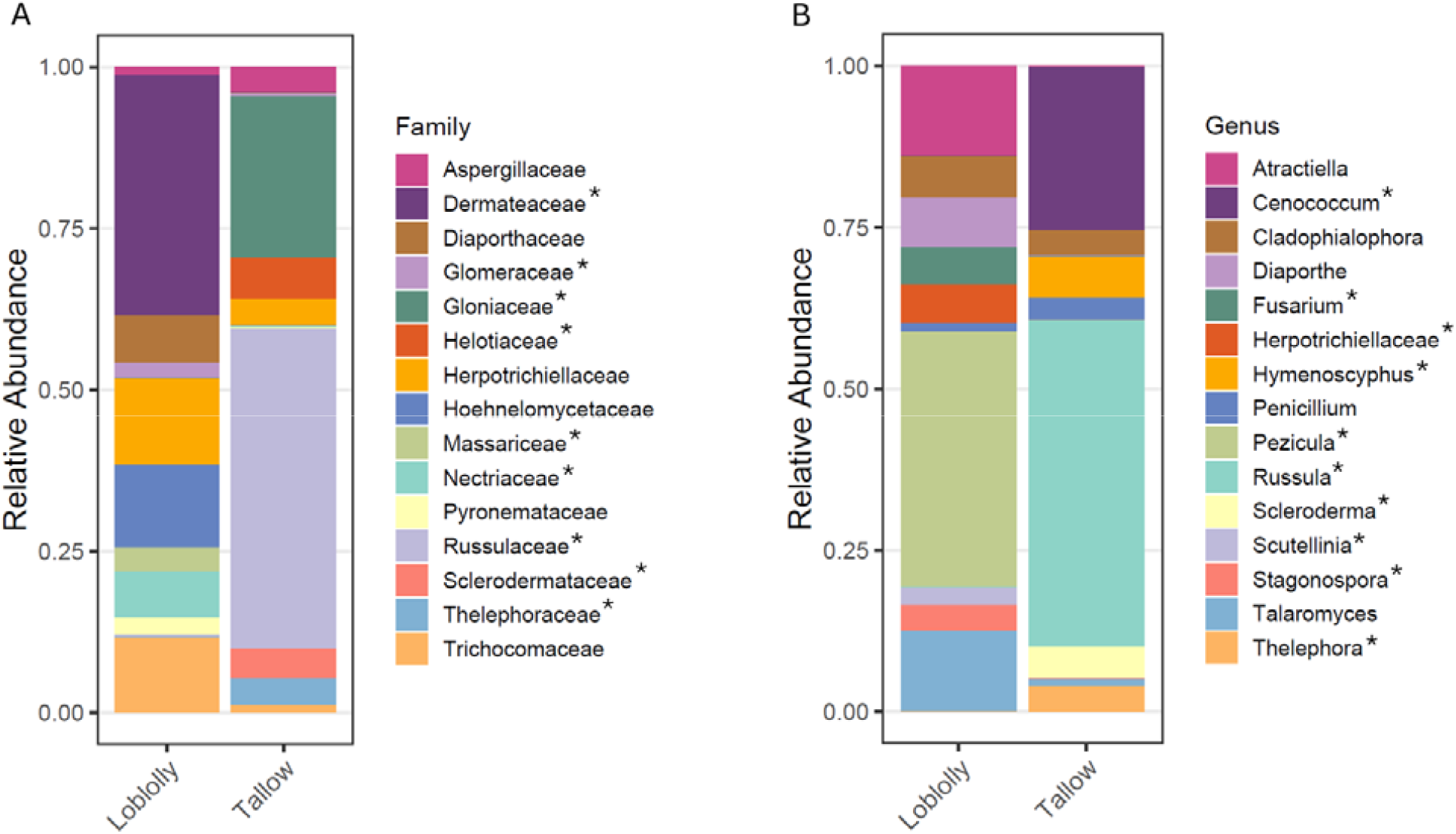
Relative abundance of the top 15 fungal families (A) and genera (B) associated with Loblolly pine and Chinese tallow rhizospheres based on ITS gene amplicon sequencing. Asterisks (*) denote taxa that differed significantly in relative abundance between Loblolly pine and Chinese tallow rhizospheres (*P* < 0.05). Each treatment consisted of three independent biological replicates.

### 3.3. Co-occurrence network analysis of bacterial and fungal communities

Both bacterial and fungal co-occurrence networks consisted of highly connected hub taxa identified based on network centrality metrics, representing the most connected members of the microbial communities (Figures 4 and 5). The bacterial co-occurrence network comprised hub taxa assigned to the genera including *Nordella, Kitasatospora, Salinispira, Leptothrix, Massilia, Novosphingobium, Opitutus, Bauldia, Rhodoplanes, Duganella, Dyella, Hirschia, Acidocella, Ideonella, Spirochaeta, Niastella, and Bacillus* (Figure 4). In contrast, the fungal co-occurrence network included hub taxa assigned to *Hymenoscyphus, Scleroderma, Cephalotheca, Scutellinia, Saitozyma, Luellia, Sebaci, Thysanorea, Inocybe, Sebaciales, Hemimycena, Collari, Dothideomycetes, Berkeleymyces*, and *Paraboeremia* (Figure 5).

**Figure 4.**
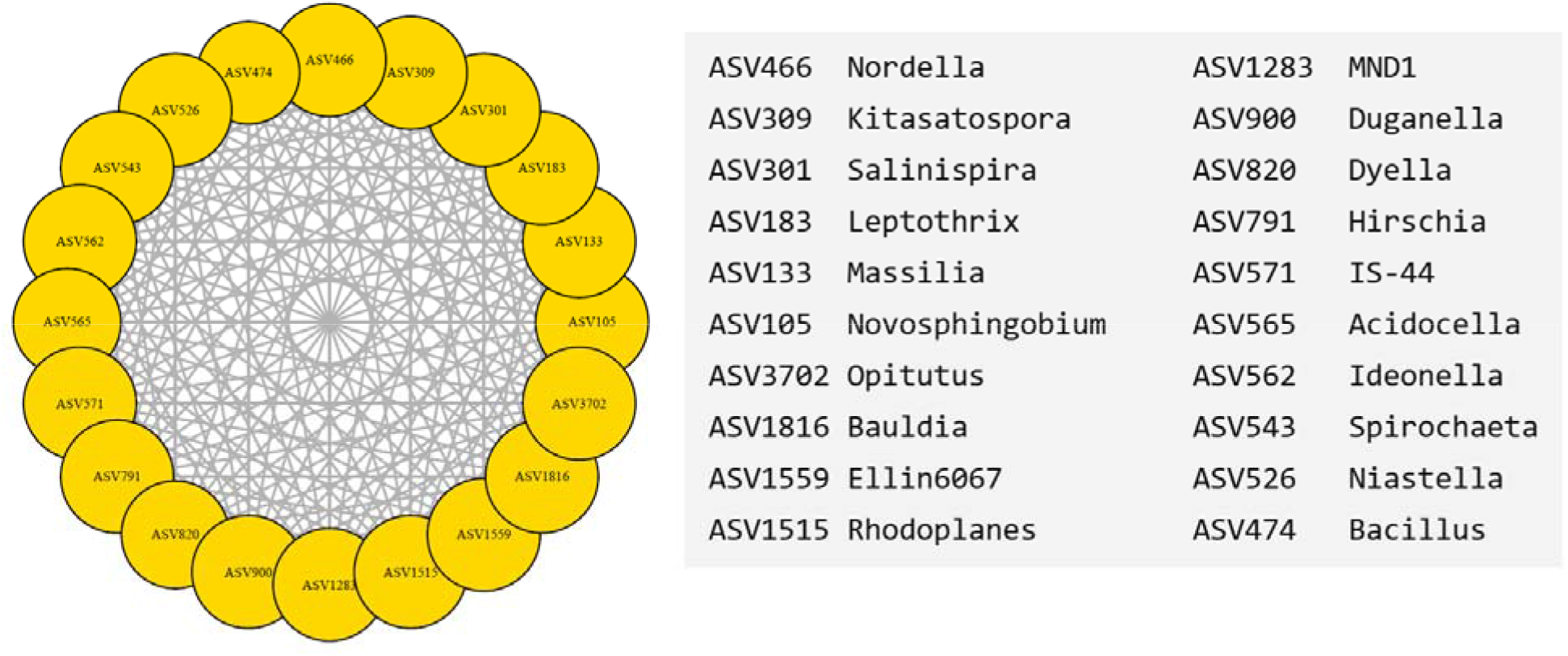
Co-occurrence network of the top 20 hub amplicon sequence variants (ASVs) identified from 16S rRNA gene amplicon sequencing of rhizosphere bacterial communities. The network illustrates the top 20 hub ASVs with the highest connectivity within the bacterial co-occurrence network. Nodes represent individual ASVs, and edges represent significant associations between ASVs. Node labels correspond to ASV identifiers. The corresponding taxonomic assignments of each hub ASV at the genus level are listed on the right. Hub taxa were identified based on network centrality metrics and represent highly connected members of the bacterial community that may play important roles in network structure and stability. Data were generated from three independent biological replicates per treatment.

**Figure 5.**
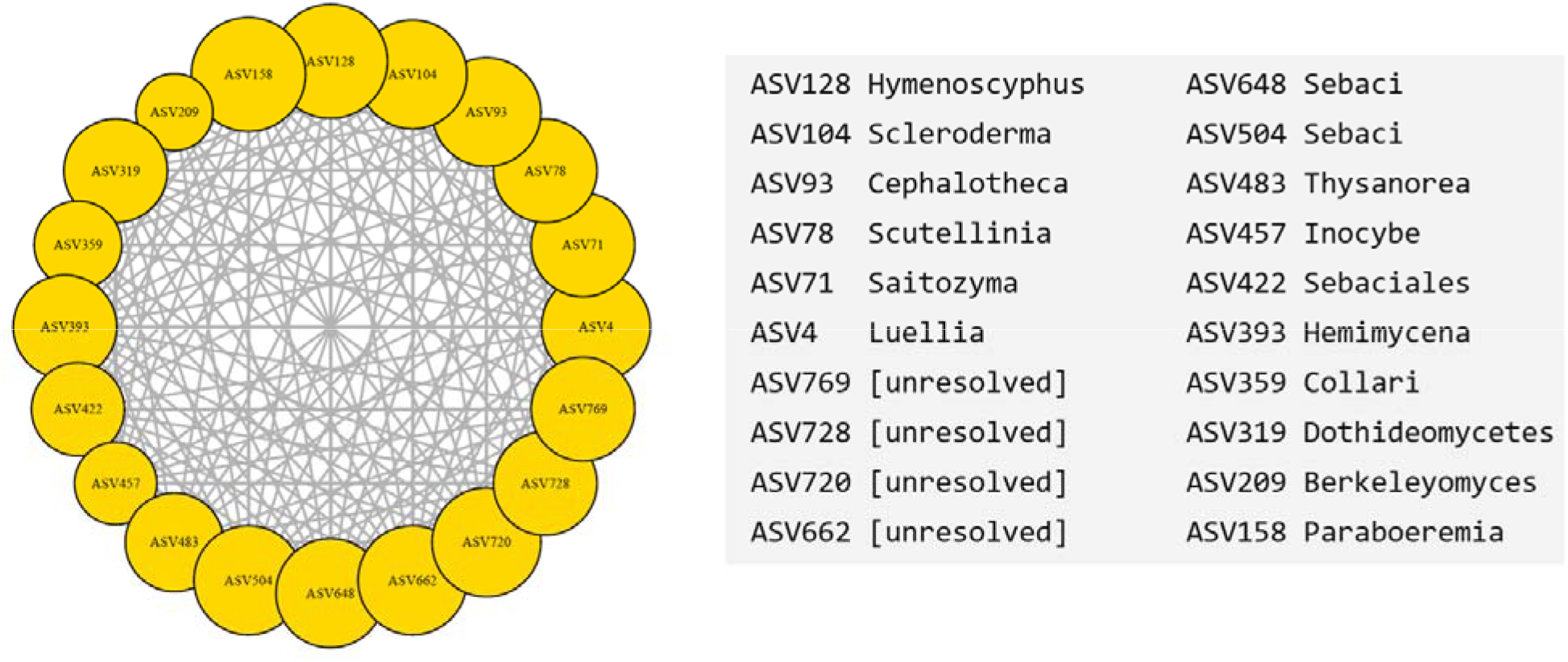
Co-occurrence network of the top 20 hub amplicon sequence variants (ASVs) identified from ITS amplicon sequencing of rhizosphere fungal communities. The network illustrates the top 20 hub ASVs with the highest connectivity within the fungal co-occurrenc network. Nodes represent individual ASVs, and edges represent significant associations between ASVs. Node labels correspond to ASV identifiers. The corresponding taxonomic assignments of each hub ASV are listed on the right. Hub taxa were identified based on network centrality metrics and represent highly connected members of the fungal community that may play important roles in network structure and stability. Data were generated from three independent biological replicates per treatment.

### 3.4. Core microbiome composition

Core microbiome analysis identified distinct sets of bacterial and fungal genera shared among samples within each treatment (Figure 6). In the bacterial community, the core microbiome of the Loblolly pine rhizosphere was dominated by *Acidothermus, Latilactobacillus, Mycobacterium, Occallatibacter*, and *Acidibacter*, whereas the Chinese tallow rhizosphere exhibited greater relative abundances of *Streptomyces, Actinoplanes, Bradyrhizobium*, and *Rhizobium*. Several core genera, including *Aquisphaera*, were detected in both rhizosphere communities but differed in their relative abundances (Figure 6A). The fungal core microbiome also differed between the two rhizosphere communities (Figure 6B). The Loblolly pine rhizosphere was characterized by relatively higher abundances of *Pezicula, Talaromyces, Cladophialophora*, and *Atractiella*, whereas the Chinese tallow rhizosphere was dominated by *Russula, Cenococcum*, and *Berkeleymyces*. Other core fungal genera, including *Hymenoscyphus* and *Diaporthe*, were present at lower relative abundances in Loblolly pine and Chinese tallow (Figure 6B).

**Figure 6.**
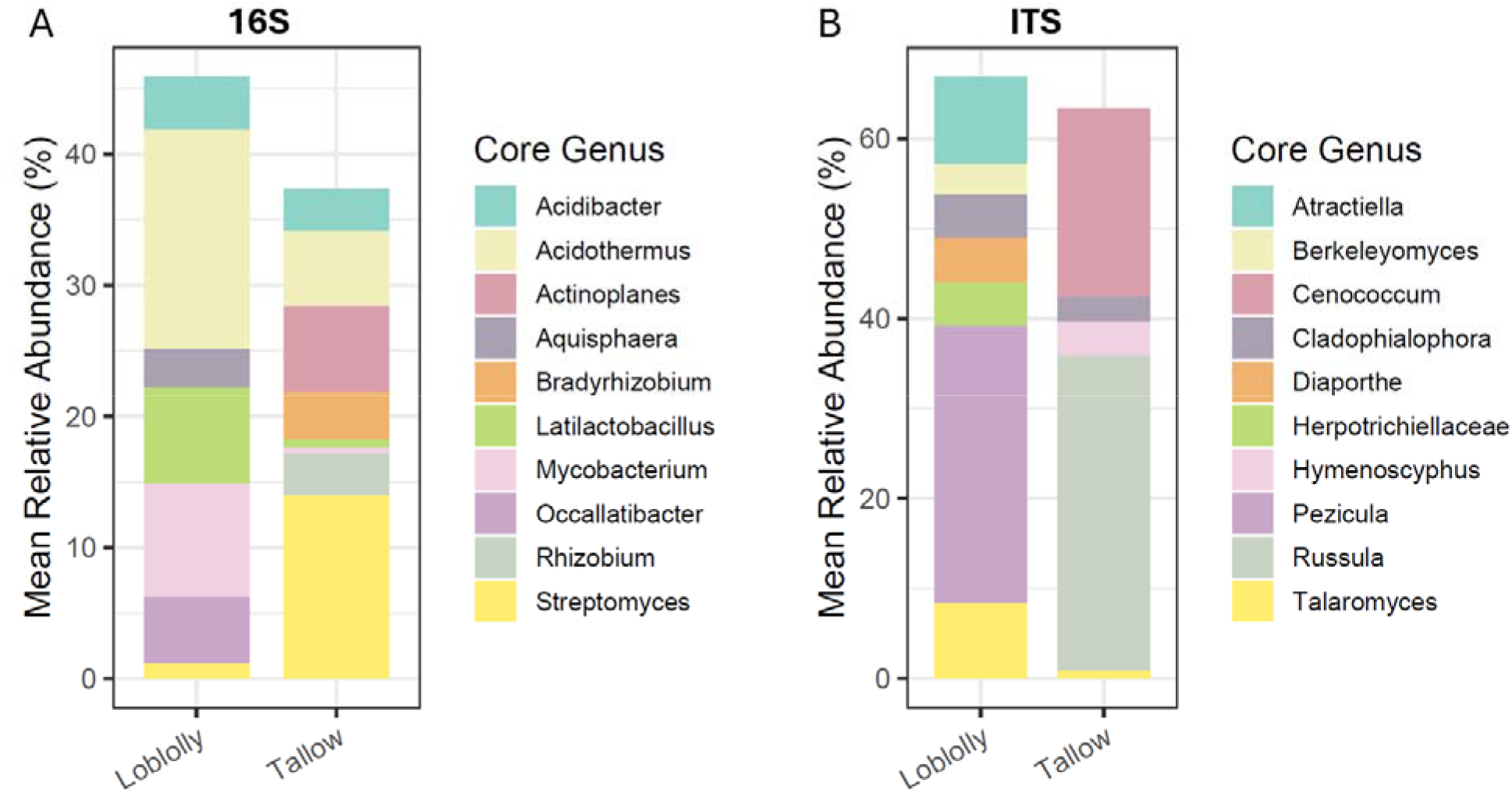
Core microbiome composition of bacterial and fungal communities associated with th rhizospheres of native loblolly pine and invasive Chinese tallow. (A) Relative abundance of th core bacterial genera identified from 16S rRNA gene amplicon sequencing. (B) Relative abundance of the core fungal genera identified from ITS amplicon sequencing. Core genera were defined as those present in ≥70% of samples within each treatment with a mean relative abundance ≥0.1%. Stacked bars represent the mean relative abundance (%) of core genera acros biological replicates.

### 3.5. Predicted functional profiles of bacterial and fungal communities

Predicted functional profiles showed that the bacterial community was primarily associated with functions related to carbohydrate degradation (cellulose, hemicellulose, and starch), which represented the most abundant predicted functional group in both rhizospheres, followed by nitrogen fixation, plant growth promotion (PGPR), antibiotic compound production, lignin degradation, nutrient mineralization, siderophore production (iron acquisition), hydrocarbon degradation, sulfur metabolism, and chitin degradation (Figure 7A). Most predicted bacterial functional groups, including carbohydrate degradation, nitrogen fixation, plant growth promotion, antibiotic compound production, nutrient mineralization, siderophore production, sulfur metabolism, and chitin degradation, exhibited higher mean relative abundances in the Chinese tallow rhizosphere than in the Loblolly pine rhizosphere. In contrast, lignin degradation and hydrocarbon degradation were relatively more abundant in the Loblolly pine rhizosphere.

**Figure 7.**
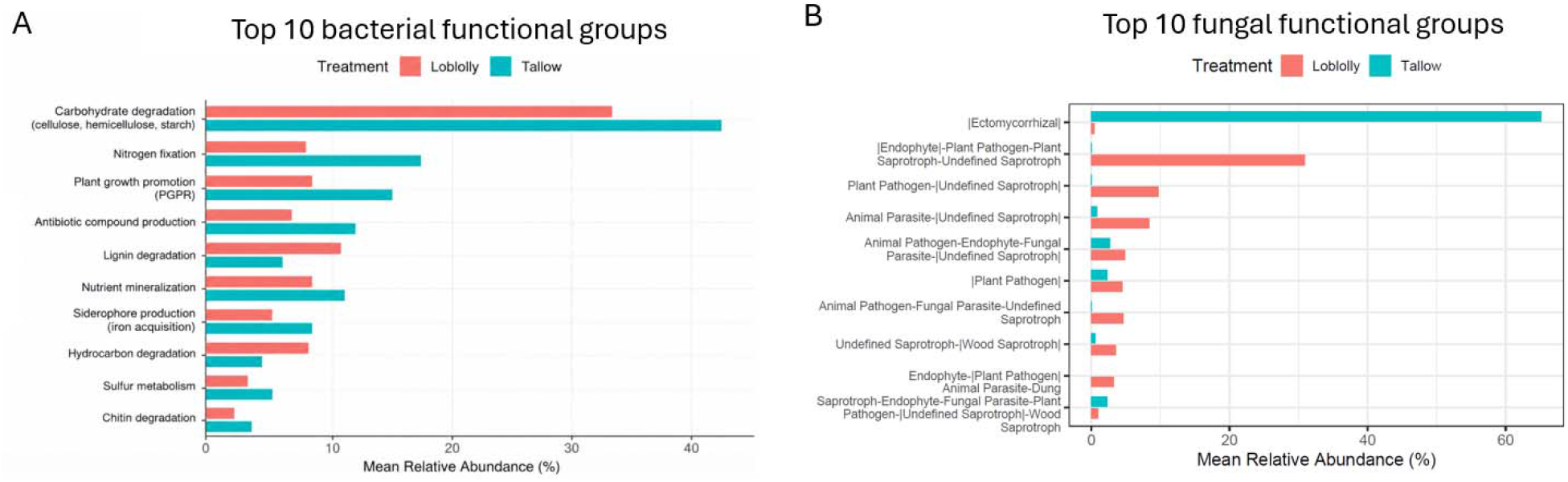
Comparison of the predicted functional profiles of fungal and bacterial communities in the rhizospheres of native loblolly pine and invasive Chinese tallow. (A). Top 10 bacterial functional groups inferred from genus-level taxonomic composition using FAPROTAX classifications and published ecological functions. (B) Top 10 fungal functional guilds predicted using FUNGuild based on ITS amplicon sequencing. Bars represent the mean relative abundance (%) of each functional group across biological replicates.

The predicted fungal functional profile also differed between the two rhizosphere communities (Figure 7B). Ectomycorrhizal fungi constituted the dominant functional guild in the Chinese tallow rhizosphere, whereas the endophyte–plant pathogen–plant saprotroph–undefined saprotroph guild was more abundant in the Loblolly pine rhizosphere. Additional guilds, including plant pathogen–undefined saprotroph, animal parasite–undefined saprotroph, animal pathogen–endophyte–fungal parasite–undefined saprotroph, plant pathogen, animal pathogen– fungal parasite–undefined saprotroph, undefined saprotroph–wood saprotroph, endophyte–plant pathogen–animal parasite–dung saprotroph, and saprotroph–endophyte–fungal parasite–plant pathogen–undefined saprotroph–wood saprotroph, occurred at relatively lower abundances in both rhizosphere communities (Figure 7B).

### 3.6. Correlations between soil physicochemical properties and dominant microbial taxa

Spearman correlation analysis revealed distinct associations between soil physicochemical properties and the dominant bacterial and fungal genera in the rhizosphere communities (Figure 8). For the bacterial community, two major clusters of taxa were observed (Figure 8A). *Mycobacterium, Latilactobacillus, Occallatibacter, Acidothermus, Conexibacter* and *Acidibacter* exhibited positive correlations with soil P, N, K, Fe, organic matter, pH, and lime buffer capacity (LBC), but negative correlations with Zn, Pb, Na, Mn, Mg, Cu, base saturation, and cation exchange capacity (CEC). In contrast, *Streptomyces, Haliangium, Bradyrhizobium, Actinoplanes, Nocardia*, and several ASVs displayed the opposite correlation pattern, showing positive associations with Zn, Pb, Na, Mn, Mg, Cu, base saturation, and CEC, and negative associations with P, N, K, Fe, organic matter, pH, and LBC (Figure 8A). The fungal community also formed two major correlation clusters (Figure 8B). *Russula, Cenococcum, Scleroderma*, and *Hymenoscyphus* were mainly positively correlated with Zn, Pb, Na, Mn, Mg, Cu, base saturation, and CEC, while exhibiting negative correlations with P, N, K, Fe, organic matter, pH, and LBC. Conversely, *Fusarium, Pezicula, Atractiella, Berkeleomyces, Scutellinia, Penicillium, Talaromyces, Cladophialophora*, and *Diaporthe* showed positive correlations with P, N, K, Fe, organic matter, pH, and LBC, but negative correlations with Zn, Pb, Na, Mn, Mg, Cu, base saturation, and CEC (Figure 8B).

**Figure 8.**
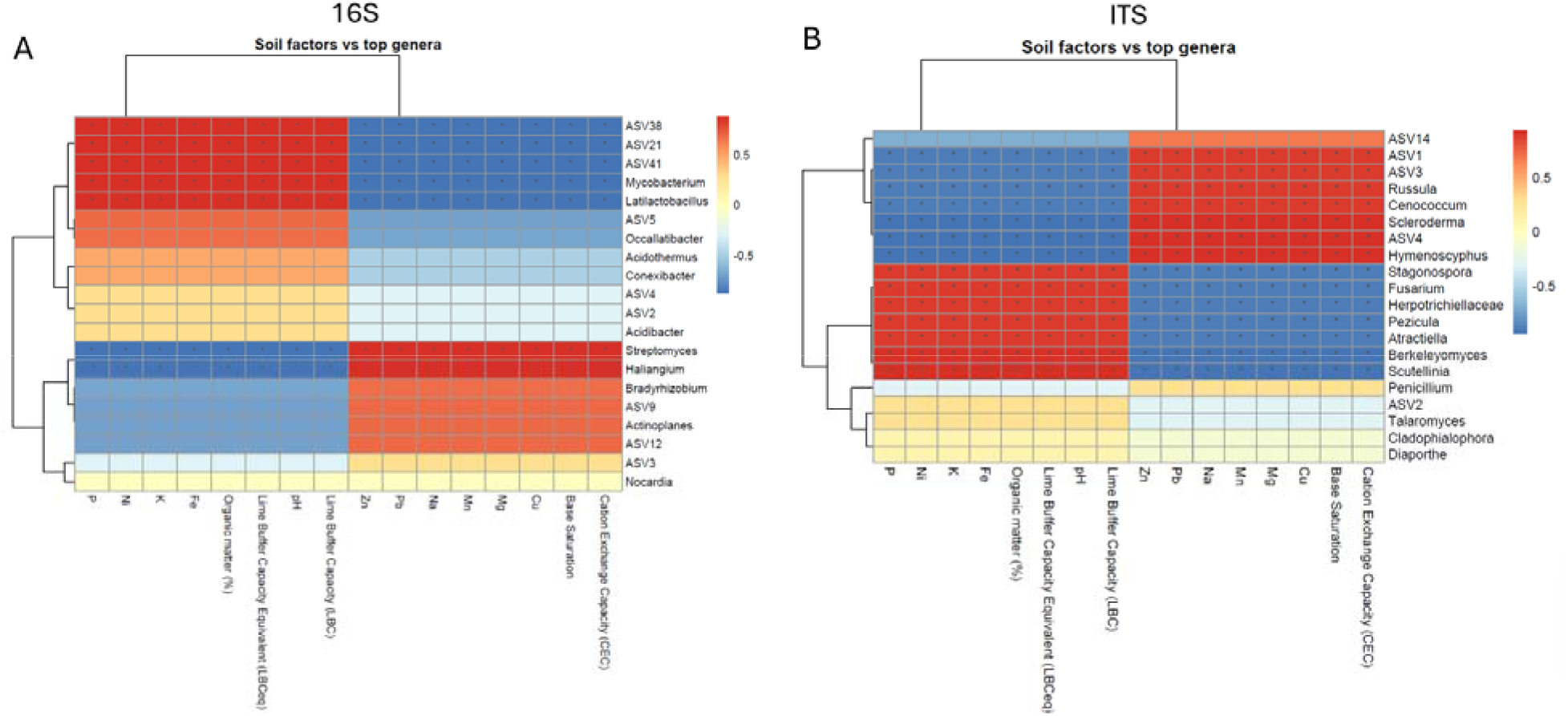
Correlations between soil physicochemical properties and dominant microbial taxa in rhizosphere communities associated with native loblolly pine and invasive Chinese tallow. (A) Heatmap showing Spearman correlation coefficients between selected soil physicochemical properties and the top bacterial genera identified from 16S rRNA gene amplicon sequencing. (B) Heatmap showing Spearman correlation coefficients between selected soil physicochemical properties and the top fungal genera identified from ITS amplicon sequencing. Rows represent microbial taxa, and columns represent measured soil properties. Color intensity indicates the strength and direction of the correlation, with red representing positive correlations and blue representing negative correlations. Hierarchical clustering was applied to group taxa and soil variables based on similarity in correlation patterns. Asterisks (*) indicate statistically significant correlations (P < 0.05). Data were generated from three independent biological replicates per treatment.

## Discussion

Interactions between plants and their rhizosphere microbiomes are increasingly recognized as important determinants of ecosystem functioning and plant adaptation to changing environments (Bouwmeester et al., 2025). In this study, we demonstrated that invasive Chinese tallow supports a rhizosphere microbiome that is taxonomically, functionally, and ecologically distinct from that of native Loblolly pine. Specifically, Chinese tallow is possibly associated with distinct bacterial and fungal communities, restructured core microbiomes and microbial networks, enriched plant-beneficial taxa and predicted nutrient-acquisition functions and altered soil–microbe associations. However, since the two host species were sampled from distinct field locations, these findings should be interpreted as host- and site-associated patterns rather than definitive host effects alone.

### 4.1. Host- and soil-associated restructuring of bacterial and fungal communities by Chinese tallow invasion

Our results indicate that Chinese tallow invasion is associated with substantial restructuring of bacterial and fungal rhizosphere communities, characterized by greater bacterial and fungal richness, distinct microbial composition, and marked shifts in dominant microbial taxa relative to the native Loblolly pine. The clear separation of bacterial and fungal communities in the PCoA analyses indicates strong host- and site-associated differences in rhizosphere microbial assembly. In addition, the significantly higher Observed ASVs and Chao1 richness in the Chinese tallow rhizosphere suggest that invasion is associated with the recruitment of a broader microbial species pool, rather than merely altering the relative abundance of existing taxa. These findings are consistent with growing evidence that invasive plants actively modify belowground microbial communities through host-specific selection rather than passively inheriting the microbial communities present in invaded soils (Bulgarelli et al., 2013; Fitzpatrick et al., 2020). Host species employ strong selective pressures on rhizosphere microorganisms through differences in nutrient uptake strategies, litter chemistry, and particularly the composition of root exudates (Rodríguez et al., 2025). Consequently, even neighboring plant species growing under similar environmental conditions frequently harbor distinct rhizosphere microbiomes (Berendsen et al., 2012; Sasse et al., 2018). Previous ecological studies have demonstrated that Chinese tallow alters litter decomposition rates, nutrient availability, and soil microbial activity, thereby generating positive plant–soil feedback that enhance its competitive dominance over native vegetation (Cameron & Spencer, 1989; Nijjer et al., 2007). The present study extends these observations by demonstrating that Chinese tallow also supports a markedly different bacterial and fungal assemblage than the co-occurring native Loblolly pine, suggesting that microbial recruitment may represent an additional belowground process associated with its invasive success.

Interestingly, differences between host species were more pronounced for community composition than for evenness. While richness estimates (Observed ASVs and Chao1) were significantly greater under Chinese tallow, Shannon and Simpson diversity did not differ significantly between treatments. This pattern suggests that invasion primarily increased the number of taxa colonizing the rhizosphere without fundamentally altering dominance relationships among abundant taxa. Similar patterns have been observed in invasive plant systems where newly recruited microorganisms occupy previously unavailable ecological niches while the dominant microbial groups remain relatively stable (Coats & Rumpho, 2014; Dawson & Schrama, 2016). Thus, Chinese tallow appears to broaden microbial recruitment rather than simply replacing dominant community members. Furthermore, the taxonomic shifts observed in both bacterial and fungal communities further support the hypothesis of host-mediated microbial filtering. Several bacterial taxa enriched in the Chinese tallow rhizosphere, including *Streptomyces, Bradyrhizobium, Rhizobium*, and *Actinoplanes*, are widely associated with nutrient acquisition, pathogen suppression, and nitrogen cycling

(Rudrappa et al., 2024; van der Meij et al., 2017; Lindström & Mousavi, 2020). Their enrichment suggests a rhizosphere environment favorable for nutrient turnover and plant growth. Although Chinese tallow is not a legume, enrichment of these genera may increase nitrogen availability within invaded soils, thereby indirectly promoting plant growth and competitive ability. Conversely, the native Loblolly pine rhizosphere showed higher abundances of *Acidothermus, Mycobacterium*, and *Latilactobacillus*. These genera were originally isolated from acidic forest soils, possesses cellulolytic capabilities that contribute to decomposition of lignocellulosic substrates characteristic of conifer litter (Bautista-Cruz et al., 2024). The greater abundance of cellulolytic taxa under Loblolly pine is therefore consistent with the chemically recalcitrant litter produced by coniferous forests, which generally favors slow-growing oligotrophic microorganisms specialized for degradation of complex carbon compounds. This observation aligns with previous studies demonstrating that litter chemistry is among the strongest determinants of forest soil microbial composition (Lladó et al., 2017; Prescott & Grayston, 2013).

The fungal community exhibited equally pronounced host specificity. Chinese tallow demonstrated significantly greater abundances of *Russula, Scleroderma, Cenococcum*, and several ectomycorrhizal-associated taxa, whereas the Loblolly pine rhizosphere contained relatively higher abundances of Pezicula, and several saprotrophic or plant pathogen-associated fungi including *Fusarium*. Although Chinese tallow has generally been considered to associate primarily with arbuscular mycorrhizal fungi, increasing evidence indicates that invasive woody plants can reshape local fungal communities by indirectly favoring ectomycorrhizal fungi through alterations in litter quality, nutrient cycling, and interactions with neighboring ectomycorrhizal hosts (Zhang et al., 2024; Dickie et al., 2017). The enrichment of ectomycorrhizal-associated genera observed here may therefore reflect invasion-induced modification of belowground ecological processes rather than obligate symbiosis with Chinese tallow itself. Furthermore, the dominance of ectomycorrhizal fungi further enhances ecosystem nutrient retention and facilitates invasion. Not only that, the observed reduction in several plant pathogen-associated fungal groups under Chinese tallow may also contribute to invasion success. Consistent with this notion, invasive plants frequently escape native soil-borne pathogens while recruiting novel or generalist mutualistic microorganisms, thereby facilitating establishment and spread (Reinhart & Callaway, 2006; Dawson & Schrama, 2016). Although the present study cannot directly determine pathogen activity, the lower relative abundance of several pathogen-associated guilds together with increased beneficial bacterial taxa is consistent with a microbial environment potentially more favorable for Chinese tallow establishment. Collectively, these findings indicate that Chinese tallow invasion substantially restructures rhizosphere microbial communities. The enrichment of beneficial microbes and decline of pathogen-associated taxa likely strengthen plant–soil feedback, promoting nutrient cycling, soil functioning, and invasion success.

### 4.2. Core microbiome reassembly, microbial networks, and predicted functional shifts

Beyond changes in microbial dynamics, our findings demonstrate that Chinese tallow invasion is associated with a substantial reorganization of the rhizosphere core microbiome and predicted functional potential. The ecological functions of rhizosphere microbiomes are governed not only by community diversity but also by the identity of persistent core taxa and the interactions among highly connected microbial populations (Chukwuneme & Babalola, 2025). Consequently, invasion-driven restructuring of the core microbiome may have greater ecological significance than changes in individual taxa because core microorganisms disproportionately influence nutrient cycling, microbial stability, and plant performance (Shade et al., 2012; Toju et al., 2018). The bacterial core microbiome exhibited clear host-specific differences between the invasive and native tree species. This distinction likely reflects differences in ecological strategies employed by the two host species. The enrichment of *Streptomyces, Bradyrhizobium*, and *Rhizobium* suggests that Chinese tallow is associated with microorganisms capable of improving nutrient availability and supporting rapid plant growth (Reinhart and Callaway, 2006). In contrast, the core bacterial community associated with Loblolly pine was dominated by taxa commonly linked with decomposition of recalcitrant organic matter characteristic of coniferous forest ecosystems. *Acidothermus*, for example, possesses highly efficient cellulolytic enzyme systems adapted to acidic forest soils and contributes to degradation of lignocellulosic substrates derived from pine litter (Barabote et al., 2009). Similarly, members of Occallatibacter and Acidibacter are frequently associated with oligotrophic forest soils where slow nutrient turnover favors microorganisms adapted to stable, carbon-limited environments (Kielak et al., 2016). These contrasting core microbiomes suggest distinct belowground nutrient acquisition strategies, with Chinese tallow favoring plant-beneficial microorganisms and Loblolly pine supporting decomposer-associated communities. The fungal core microbiome further supports this ecological shift. Chinese tallow was enriched in *Russula* and *Cenococcum*, widespread ectomycorrhizal fungi known to enhance nutrient acquisition and drought tolerance in forest ecosystems. The enrichment of ectomycorrhizal-associated fungi observed here therefore likely reflects broader invasion-driven changes in belowground ecological processes rather than exclusive host specificity.

Microbial co-occurrence network analysis identified bacterial hubs such as *Massilia, Novosphingobium*, and *Bacillus*, together with fungal hubs including *Scleroderma, Inocybe*, and members of *Sebacinales*. These taxa are recognized as keystone members of forest rhizosphere microbiomes because of their roles in nutrient cycling, pathogen suppression, and mycorrhizal functioning (Dai et al., 2025; Zhang et al., 2024; Wang et al., 2023). Although our network analyses do not establish direct ecological interactions, they demonstrate that invasion restructures highly connected microbial populations that may disproportionately influence community resilience and ecosystem functioning. Predicted analysis also suggested that bacterial communities associated with Chinese tallow possessed greater representation of functions related to carbohydrate degradation, nitrogen fixation, plant growth promotion, nutrient mineralization, and siderophore production, whereas lignin degradation and hydrocarbon degradation were relatively more prominent in the Loblolly pine rhizosphere. Although functional prediction from marker-gene sequencing cannot replace metagenomic analyses, these patterns are biologically consistent with the observed taxonomic composition. Such functional traits are expected to accelerate nutrient turnover and increase resource availability in invaded soils, potentially reinforcing the rapid growth strategy characteristic of Chinese tallow. Similarly, FUNGuild analysis indicated a greater relative abundance of ectomycorrhizal guilds in the Chinese tallow rhizosphere, whereas Loblolly pine retained higher proportions of saprotrophic and plant pathogen-associated guilds. Mycorrhizal fungi enhance nutrient foraging efficiency and improve host tolerance to multiple environmental stresses, while excessive accumulation of plant pathogens can limit plant performance (Bever et al., 2010). Taken together, our findings indicate that Chinese tallow invasion is associated with coordinated reassembly of the rhizosphere microbiome, including shifts in core taxa, microbial networks, and predicted functional potential. These changes likely reinforce nutrient acquisition, ecosystem functioning, and positive soil biotic feedbacks, thereby contributing to the invasion success of Chinese tallow.

### 4.3. Soil physicochemical filtering links microbial assembly to ecosystem functioning and invasion success

Rhizosphere microbial communities are assembled through the combined influences of host plants and the surrounding soil environment (Han et al., 2025). Our correlation analyses indicate that soil nutrients and chemical properties were closely associated with the distribution of dominant bacterial and fungal taxa, suggesting that environmental filtering acts together with host-mediated selection to shape microbial community assembly during Chinese tallow invasion. These findings support the view that rhizosphere microbiomes are shaped by both host selection and environmental filtering (Fierer, 2017; Trivedi et al., 2020). Chinese tallow-associated soils exhibited higher exchangeable base cations and cation exchange capacity, whereas Loblolly pine-associated soils contained greater iron, phosphorus, and organic matter. High cation exchange capacity generally increases the retention of essential mineral nutrients within soil, thereby enhancing nutrient availability for both microorganisms and plants (Nel et al., 2023). The enrichment of plant-beneficial microorganisms under these soil conditions suggests positive feedback in which invasion-induced modification of soil chemistry may favor microbial groups capable of accelerating nutrient cycling (Dawson and Schrama, 2016). In contrast, bacterial genera enriched in the Loblolly pine rhizosphere are commonly associated with forest ecosystems characterized by slower nutrient turnover and greater dependence on decomposition of structurally complex organic matter. Coniferous forests typically accumulate litter enriched in lignin, cellulose, and phenolic compounds that decompose relatively slowly, selecting for microorganisms capable of degrading recalcitrant carbon substrates (Pennanen et al., 1999). The stronger representation of cellulolytic bacteria together with higher predicted lignocellulose degradation observed in Loblolly pine therefore appears consistent with the conservative nutrient cycling strategy characteristic of many pine-dominated ecosystems. In addition, the fungal community exhibited similarly strong relationships with soil chemistry. Ectomycorrhizal-associated genera were positively associated with higher cation exchange capacity and base saturation, whereas genera including *Pezicula, Cladophialophora, Fusarium*, and *Talaromyces* showed stronger associations with soils containing greater organic matter and iron. These patterns further support the hypothesis that invasion restructures fungal community assembly through modification of soil nutrient availability. Mycorrhizal fungi contribute substantially to phosphorus mobilization, nitrogen acquisition, and water uptake while simultaneously influencing soil aggregation and carbon sequestration (Huey et al., 2020).

An important implication of our findings is that microbial restructuring may not simply reflect invasion outcomes; instead, microorganisms may participate in reciprocal soil biotic feedbacks that reinforce ecosystem change. Chinese tallow has previously been shown to alter litter decomposition, nutrient mineralization, and microbial respiration following establishment (Cameron and Spencer, 1989; Nijjer et al., 2007). Our data suggest that these ecosystem changes are accompanied by coordinated shifts in microbial composition, interaction networks, and predicted functional traits that collectively favor microorganisms associated with rapid nutrient cycling and plant growth promotion. Such rhizosphere-mediated interactions enhance stress tolerance while promoting long-term invasion success (Bever et al., 2010). Overall, our findings support a model in which Chinese tallow invasion restructures rhizosphere microbial communities through coordinated host-mediated selection and changes in soil physicochemical properties (Figure 9). Also, the belowground changes likely promote nutrient acquisition, ecosystem functioning, and beneficial plant–microbe interactions, facilitating the continued establishment and spread of Chinese tallow in southeastern U.S. forests. Although FAPROTAX and FUNGuild provide useful functional predictions, they infer function from taxonomy rather than directly measuring microbial activity. Future studies integrating multi-omics approaches and experimental microbiome manipulation are needed to validate these predicted functions and establish causal links between rhizosphere microbiomes and Chinese tallow invasion.

**Figure 9.**
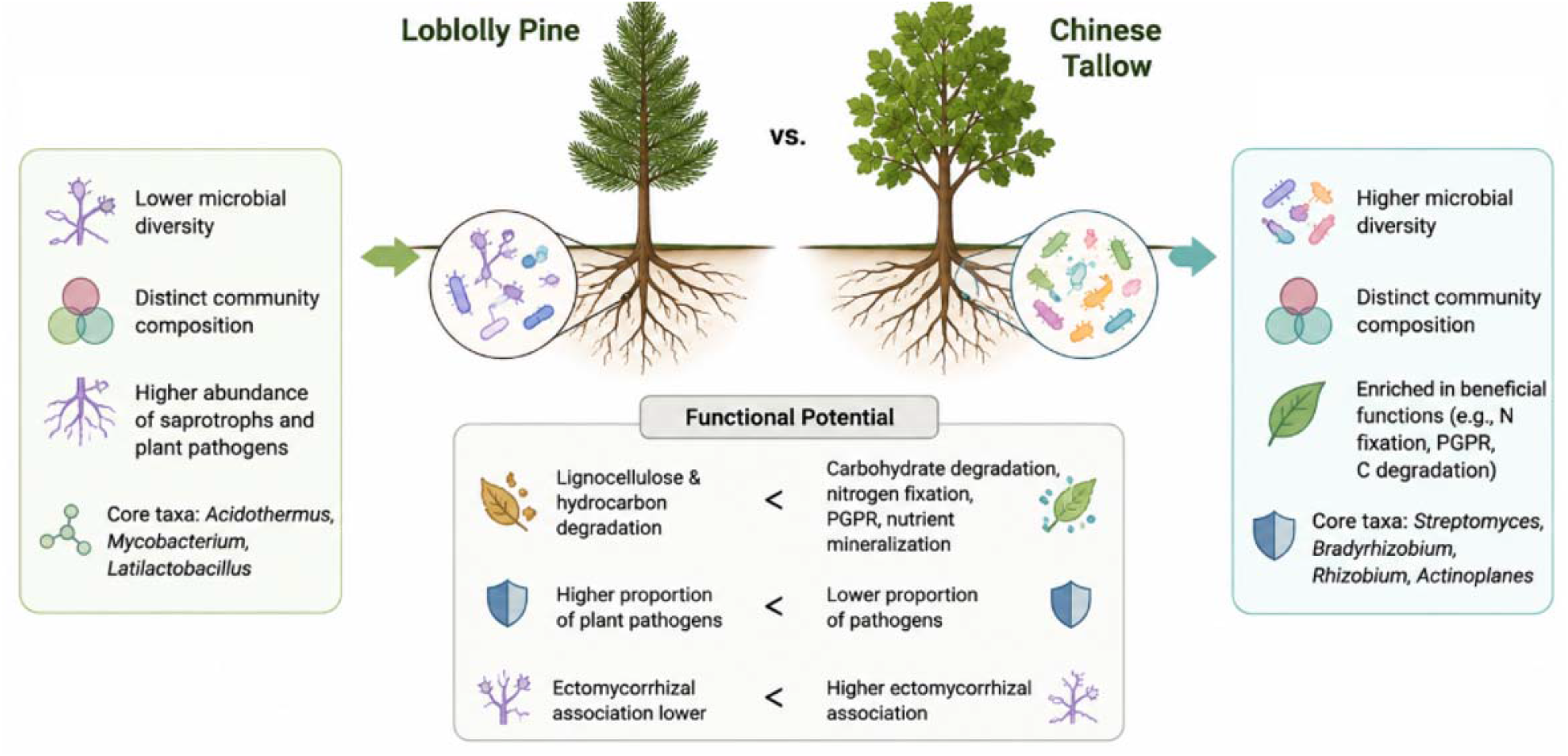
Conceptual model illustrating the contrasting rhizosphere microbial communities and predicted functional potential associated with native loblolly pine and invasive Chinese tallow. Chinese tallow supports greater microbial diversity, distinct community composition, enrichment of beneficial bacterial functions (e.g., nitrogen fixation, plant growth promotion, and carbohydrate degradation), and increased ectomycorrhizal fungi, whereas loblolly pine is associated with greater lignocellulose degradation and relatively higher abundances of saprotrophic and plant pathogen-associated fungi. These host-specific microbial shifts may influence nutrient cycling and ecosystem functioning.

## Conclusion

Our study demonstrates that Chinese tallow invasion is associated with a fundamental reorganization of rhizosphere microbial communities, extending beyond changes in diversity to the assembly of distinct bacterial and fungal consortia with contrasting ecological characteristics. The enrichment of plant-beneficial bacterial taxa (*Streptomyces, Bradyrhizobium*, and *Rhizobium*) and ectomycorrhizal-associated fungi (*Russula* and *Cenococcum*), together with the decline of taxa associated with conservative nutrient cycling in native Loblolly pine, suggests a shift toward a rhizosphere environment that favors rapid nutrient acquisition and plant performance. By integrating bacterial and fungal community analyses with soil physicochemical properties, this study provides a comprehensive perspective on the belowground ecological changes associated with woody plant invasion. Given the field-based design, future reciprocal soil inoculation and multi-site studies are needed to separate host-specific effects from site-level environmental filtering. These findings indicate that rhizosphere microbiome reassembly is a key ecological feature of Chinese tallow invasion and underscore the value of incorporating microbial communities into invasion ecology, forest management, and restoration strategies aimed at improving the resilience of invaded ecosystems.

## Acknowledgements

The authors thank LC Sciences (Houston, TX, USA) for performing the 16S rRNA gene and ITS amplicon sequencing and for providing high-quality sequencing data. We also acknowledge the technical support provided throughout the sequencing process.

## Data availability statement

The raw amplicon sequencing data generated in this study are publicly available in the NCBI BioProject database under accession number PRJNA1482808 (https://www.ncbi.nlm.nih.gov/bioproject/).

## References

Barabote, R. D., Xie, G., Leu, D. H., Normand, P., Necsulea, A., Daubin, V., Médigue, C., Adney, W. S., Xu, X. C., Lapidus, A., Parales, R. E., Detter, C., Pujic, P., Bruce, D., Lavire, C., Challacombe, J. F., Brettin, T. S., & Berry, A. M. (2009). Complete genome of the cellulolytic thermophile Acidothermus cellulolyticus 11B provides insights into its ecophysiological and evolutionary adaptations. Genome research, 19(6), 1033–1043.

Bardgett, R. D., & van der Putten, W. H. (2014). Belowground biodiversity and ecosystem functioning. Nature, 515(7528), 505–511. 10.1038/nature13855

Bautista-Cruz, A., Aquino-Bolaños, T., Hernández-Canseco, J., & Quiñones-Aguilar, E. E. (2024). Cellulolytic Aerobic Bacteria Isolated from Agricultural and Forest Soils: An Overview. Biology, 13(2), 102.

Berendsen, R. L., Pieterse, C. M., & Bakker, P. A. (2012). The rhizosphere microbiome and plant health. Trends in plant science, 17(8), 478–486.

Bever, J. D., Platt, T. G., & Morton, E. R. (2012). Microbial population and community dynamics on plant roots and their feedbacks on plant communities. Annual review of microbiology, 66, 265–283.

Bonito, G., Smith, M. E., Nowak, M., Healy, R. A., Guevara, G., Cázares, E., Kinoshita, A., Nouhra, E. R., Domínguez, L. S., Tedersoo, L., Murat, C., Wang, Y., Moreno, B. A., Pfister, D. H., Nara, K., Zambonelli, A., Trappe, J. M., & Vilgalys, R. (2013). Historical biogeography and diversification of truffles in the Tuberaceae and their newly identified southern hemisphere sister lineage. PloS one, 8(1), e52765.

Bouwmeester, H., Dong, L., Wippel, K., Hofland, T., & Smilde, A. (2025). The chemical interaction between plants and the rhizosphere microbiome. Trends in plant science, 30(9), 1002–1019.

Brar, B., Bala, K., Saharan, B. S., Sadh, P. K., & Duhan, J. S. (2024). Bio-boosting agriculture: Harnessing the potential of fungi-bacteria-plant synergies for crop improvement. Discover Plants, 1, Article 21. 10.1007/s44372-024-00023-0

Bruce, K. A., Cameron, G. N., Harcombe, P. A., & Jubinsky, G. (1997). Introduction, impact on native habitats, and management of a woody invader, the Chinese tallow tree (Sapium sebiferum (L.) Roxb.). Natural Areas Journal, 17(3), 255–260.

Bulgarelli, D., Schlaeppi, K., Spaepen, S., Ver Loren van Themaat, E., & Schulze-Lefert, P. (2013). Structure and functions of the bacterial microbiota of plants. Annual review of plant biology, 64, 807–838.

Callahan, B. J., McMurdie, P. J., Rosen, M. J., Han, A. W., Johnson, A. J. & Holmes, S. P. (2016) DADA2: High-resolution sample inference from Illumina amplicon data. Nature Methods, 13(7), 581–583.

Cameron, G. N., & Spencer, S. R. (1989). Rapid leaf decay and nutrient release in a chinese tallow forest. Oecologia, 80(2), 222–228.

Chukwuneme, C. F., & Babalola, O. O. (2025). Microbial diversity and function in the rhizosphere microbiome: Driving forces and monitoring approaches. Agrosystems, Geosciences & Environment, 8(3), e70169. 10.1002/agg2.70169

Clarke J. D. (2009). Cetyltrimethyl ammonium bromide (CTAB) DNA miniprep for plant DNA isolation. Cold Spring Harbor protocols, 2009(3), pdb.prot5177.

Coats, V. C., & Rumpho, M. E. (2014). The rhizosphere microbiota of plant invaders: an overview of recent advances in the microbiomics of invasive plants. Frontiers in microbiology, 5,368.

Compant, S., van der Heijden, M. G., & Sessitsch, A. (2010). Climate change effects on beneficial plant-microorganism interactions. FEMS microbiology ecology, 73(2), 197–214.

Dai, X., Yang, P., Zhou, C., Chen, Z., Li, S., & Zhu, T. (2025). Bacillus mojavensis dxk33 Modulates Rhizosphere Microbiome and Suppresses Root Rot in Cunninghamia lanceolata. Microorganisms, 14(1), 34.

Dawson, W., & Schrama, M. (2016). Identifying the role of soil microbes in plant invasions. Journal of Ecology, 104(5), 1211–1218. 10.1111/1365-2745.12619

Dickie, I. A., Bufford, J. L., Cobb, R. C., Desprez-Loustau, M. L., Grelet, G., Hulme, P. E., Klironomos, J., Makiola, A., Nuñez, M. A., Pringle, A., Thrall, P. H., Tourtellot, S. G., Waller, L., & Williams, N. M. (2017). The emerging science of linked plant-fungal invasions. The New phytologist, 215(4), 1314–1332.

Fierer, N., & Jackson, R. B. (2006). The diversity and biogeography of soil bacterial communities. Proceedings of the National Academy of Sciences of the United States of America, 103(3), 626–631.

Fierer, N., Wood, S. A., & Bueno de Mesquita, C. P. (2021). How microbes can, and cannot, be used to assess soil health. Soil Biology and Biochemistry, 153, 108111. 10.1016/j.soilbio.2020.108111

Fitzpatrick, C. R., Salas-González, I., Conway, J. M., Finkel, O. M., Gilbert, S., Russ, D., Teixeira, P. J. P. L., & Dangl, J. L. (2020). The Plant Microbiome: From Ecology to Reductionism and Beyond. Annual review of microbiology, 74, 81–100.

Han, M., Han, Y., Liu, X., Li, G., & Li, P. (2025). Rhizosphere microbiomes altered by environmental stresses and agronomic practices: Implications for plant adaptation and soil biogeochemical processes. Plant Stress, 18, 101062.

Han, X., Xia, T., Shen, K., Wu, B., & He, Y. (2026). Effects of arbuscular mycorrhizal fungus on the functional bacteria assembly in promoting plant growth and nutrient acquisition in karst soil. BMC microbiology, 26(1), 196.

Harrell FEJ (2012) Package ‘Hmisc’ – Harrell Miscellaneous. http://cranr-projectorg/web/packages/Hmisc/Hmiscpdf: 1–362.

Hodgson, R. J., Cando-Dumancela, C., Liddicoat, C., Ramesh, S. A., Edwards, R. A., & Breed, M. F. (2025). Strong Host Modulation of Rhizosphere-to-Endosphere Microbial Colonisation in Natural Populations of the Pan-Palaeotropical Keystone Grass Species, Themeda triandra. Ecology and evolution, 15(6), e71595.

Huey, C. J., Gopinath, S. C. B., Uda, M. N. A., Zulhaimi, H. I., Jaafar, M. N., Kasim, F. H., & Yaakub, A. R. W. (2020). Mycorrhiza: a natural resource assists plant growth under varied soil conditions. 3 Biotech, 10(5), 204.

Kielak, A. M., Barreto, C. C., Kowalchuk, G. A., van Veen, J. A., & Kuramae, E. E. (2016). The Ecology of Acidobacteria: Moving beyond Genes and Genomes. Frontiers in microbiology, 7, 744.

Lindström, K., & Mousavi, S. A. (2020). Effectiveness of nitrogen fixation in rhizobia. Microbial biotechnology, 13(5), 1314–1335.

Lladó, S., López-Mondéjar, R., & Baldrian, P. (2017). Forest Soil Bacteria: Diversity, Involvement in Ecosystem Processes, and Response to Global Change. Microbiology and molecular biology reviews : MMBR, 81(2), e00063–16.

Louca S., Parfrey L.W., Doebeli M. Decoupling function and taxonomy in the global ocean microbiome. Science. 2016;353:1272–1277.

Lowry, C. J., Blocklove, K. M., Eckert, K. E., Koirala, N., & Rice, E. K. (2024). If you can’t beat them, join them: Using invasive plants to understand rhizodeposits as drivers of plant–soil feedbacks. Frontiers in Agronomy, 6, Article 1363124. 10.3389/fagro.2024.1363124

Lustenhouwer, N., Chaubet, T. M. R., Melen, M. K., van der Putten, W. H., & Parker, I. M. (2024). Plant-soil interactions during the native and exotic range expansion of an annual plant. Journal of evolutionary biology, 37(6), 653–664.

McMurdie, P. J., & Holmes, S. (2013). phyloseq: an R package for reproducible interactive analysis and graphics of microbiome census data. PloS one, 8(4), e61217.

Meng, F., Duan, R., Yang, H., Dai, Q., Zhang, Y., & Fu, J. (2025). Advances in Plant Species Recognition Mediated by Root Exudates: A Review. Plants (Basel, Switzerland), 14(19), 3076.

Natural Resources Conservation Service (NRCS) (2008). General soil map of Texas. U.S. Department of Agriculture. https://www.nrcs.usda.gov/sites/default/files/2023-08/Texas%20General%20Soil%20Map.pdf

Natural Resources Conservation Service (NRCS). 2026. Web Soil Survey. U.S. Department of Agriculture. https://websoilsurvey.nrcs.usda.gov

Nel, T., Bruneel, Y., & Smolders, E. (2023). Comparison of five methods to determine the cation exchange capacity of soil. Journal of Plant Nutrition and Soil Science, 186(4), 459–469. 10.1002/jpln.202200378

Nguyen NH, Song Z, Bates ST, et al., FUNGuild: an open annotation tool for parsing fungal community datasets by ecological guild. Fungal Ecol. 2016;20:241–248.

Nijjer, S., Rogers, W. E., & Siemann, E. (2007). Negative plant-soil feedbacks may limit persistence of an invasive tree due to rapid accumulation of soil pathogens. Proceedings. Biological sciences, 274(1625), 2621–2627.

Nilsson, R. H., Larsson, K. H., Taylor, A. F. S., Bengtsson-Palme, J., Jeppesen, T. S., Schigel, D., Kennedy, P., Picard, K., Glöckner, F. O., Tedersoo, L., Saar, I., Kõljalg, U., & Abarenkov, K. (2019). The UNITE database for molecular identification of fungi: handling dark taxa and parallel taxonomic classifications. Nucleic acids research, 47(D1), D259–D264.

Pechlivanis, N., Karakatsoulis, G., Kyritsis, K., Tsagiopoulou, M., Sgardelis, S., Kappas, I., & Psomopoulos, F. (2024). Microbial co-occurrence network demonstrates spatial and climatic trends for global soil diversity. Scientific data, 11(1), 672.

Pennanen, T., Liski, J., Bååth, E., Kitunen, V. V, Uotila, J., Westman, C. J., & Fritze, H. (1999). Structure of the Microbial Communities in Coniferous Forest Soils in Relation to Site Fertility and Stand Development Stage. Microbial ecology, 38(2), 168–179.

Philippot, L., Raaijmakers, J. M., Lemanceau, P., & van der Putten, W. H. (2013). Going back to the roots: The microbial ecology of the rhizosphere. Nature Reviews Microbiology, 11(11), 789–799. 10.1038/nrmicro3109

Pile, L. S., Wang, G. G., Waldrop, T. A., Walker, J. L., Bridges, W. C., & Layton, P. A. (2017). Managing an established tree invader: Developing control methods for Chinese tallow (Triadica sebifera) in maritime forests. Journal of Forestry, 115(6), 522–529.

Prescott, C. E., & Grayston, S. J. (2013). Tree species influence on microbial communities in litter and soil: Current knowledge and research needs. Forest Ecology and Management, 309, 19–27.

Quast, C., Pruesse, E., Yilmaz, P., Gerken, J., Schweer, T., Yarza, P., Peplies, J., & Glöckner, F. O. (2013). The SILVA ribosomal RNA gene database project: improved data processing and web-based tools. Nucleic acids research, 41(Database issue), D590–D596.

Reinhart, K. O., & Callaway, R. M. (2006). Soil biota and invasive plants. The New phytologist, 170(3), 445–457.

Rodríguez, M.N., Pioppi, A., & Kovács, Á. T. (2025). The role of plant host genetics in shaping the composition and functionality of rhizosphere microbiomes. mSystems, 10(8), e0004124.

Rout, M. E., & Callaway, R. M. (2012). Interactions between exotic invasive plants and soil microbes in the rhizosphere suggest that ‘everything is not everywhere’. Annals of botany, 110(2), 213–222.

Rudrappa, M., Santosh Kumar, M., Basavarajappa, D. S., Hiremath, H., Hugar, A., Almansour, A. I., Kantli, G. B., & Nayaka, S. (2024). Bioproduction, purification, partial characterization and phenol removal efficacy of tyrosinase enzyme from Streptomyces sp. strain MR28. Environmental research, 251(Pt 2), 118701.

Sasse, J., Martinoia, E., & Northen, T. (2018). Feed Your Friends: Do Plant Exudates Shape the Root Microbiome?. Trends in plant science, 23(1), 25–41.

Shade, A., Peter, H., Allison, S. D., Baho, D. L., Berga, M., Bürgmann, H., Huber, D. H., Langenheder, S., Lennon, J. T., Martiny, J. B., Matulich, K. L., Schmidt, T. M., & Handelsman, J. (2012). Fundamentals of microbial community resistance and resilience. Frontiers in microbiology, 3, 417.

Toju, H., Peay, K. G., Yamamichi, M., Narisawa, K., Hiruma, K., Naito, K., Fukuda, S., Ushio, M., Nakaoka, S., Onoda, Y., Yoshida, K., Schlaeppi, K., Bai, Y., Sugiura, R., Ichihashi, Y., Minamisawa, K., & Kiers, E. T. (2018). Core microbiomes for sustainable agroecosystems. Nature plants, 4(5), 247–257.

Trivedi, P., Leach, J. E., Tringe, S. G., Sa, T., & Singh, B. K. (2020). Plant-microbiome interactions: from community assembly to plant health. Nature reviews. Microbiology, 18(11), 607–621.

van der Meij, A., Worsley, S. F., Hutchings, M. I., & van Wezel, G. P. (2017). Chemical ecology of antibiotic production by actinomycetes. FEMS microbiology reviews, 41(3), 392–416.

Wang, H. H., Grant, W. E., Gan, J., Rogers, W. E., Swannack, T. M., Koralewski, T. E., Miller, J. H., & Taylor, J. W. (2012). Integrating spread dynamics and economics of timber production to manage Chinese tallow invasions in southern U.S. forestlands. PloS one, 7(3), e33877.

Wang, H. R., Du, X. R., Zhang, Z. Y., Feng, F. J., & Zhang, J. M. (2023). Rhizosphere interface microbiome reassembly by arbuscular mycorrhizal fungi weakens cadmium migration dynamics. iMeta, 2(4), e133.

Wang, R., Wang, Z., Liao, W., Wang, T., & Su, Y. (2026). Mikania micrantha invasion restructures rhizosphere nitrogen cycling through enzyme activation, microbial recruitment, and allelopathic regulation. Microbiome, 14(1), 97.

Yue, H., Yue, W., Jiao, S., Kim, H., Lee, Y. H., Wei, G., Song, W., & Shu, D. (2023). Plant domestication shapes rhizosphere microbiome assembly and metabolic functions. Microbiome, 11(1), 70.

Zhang, W., Xue, W., Liu, J., Zhu, H., & Zhao, Z. (2024). Molecular Diversity of Ectomycorrhizal Fungi in Relation to the Diversity of Neighboring Plant Species. Microorganisms, 12(8), 1718.

